# Effects of Livestock Grazing Exclusion on Perennial Grasses and Woody Species on the San Pedro Riparian National Conservation Area, Arizona

**DOI:** 10.1101/2022.06.23.497368

**Authors:** Marcia F. Radke

## Abstract

The objective of this study was to determine the effects of livestock exclusion on native and non-native perennial grass richness and frequency, and woody species frequency on the San Pedro Riparian National Conservation Area. A significant increase in species richness of native perennial grass species was noted overall and in retired agricultural fields, supporting the hypothesis of increased species richness with grazing exclusion. There was a significant increase in the number of plots that contained an increase in bristlegrass (*Setaria macrostachya* Kunth and *S. leucopila* (Scribn. & Merr.) K. Schum.). No significant change in species richness of non-native invasive perennial grass species was observed. Lehmann lovegrass (*Eragrostis lehmanniana* Nees) had a significant increase in the number of plots that contained the species, supporting the hypothesis that Lehmann lovegrass expansion would occur in previously uninvaded areas even without current livestock grazing. Velvet mesquite (*Prosopis velutina* Wooton) exhibited a significant increase in canopy frequency, but velvet mesquite basal frequency showed no significant increase or decrease. Results indicate that past grazing was a primary factor leading to the current distribution and density of mesquite on the SPRNCA because the same basal (i.e. rooted stem) frequency existed prior to and after grazing exclusion. Management considerations include continued grazing exclusion, inventory and monitoring of pollinator plants, continued monitoring of pace frequency transects, use of prescribed fire in uplands, monitoring of soil erosion, and implementation of erosion controls.

## INTRODUCTION

The San Pedro Riparian National Conservation Area (SPRNCA) was established in 1988 by Public Law 100-696 in a manner that “conserves, protects, and enhances the riparian area” including the aquatic and wildlife resources therein, and is managed by the U.S. Bureau of Land Management (BLM). As part of historic Spanish land grants, the SPRNCA has long been grazed by livestock. However, a grazing moratorium on the SPRNCA went into effect in 1989 (54 FR 38058), which remains in effect. Pace frequency transects on SPRNCA were established in 1987 and monitored to determine the effects of grazing exclusion on wildlife habitat.

Native species richness was reduced by increasing grazing intensity at all spatial scales (Dorrough et al. 2007), or, if grazing intensifies beyond the evolutionary history, diversity is predicted to decline sharply and richness will accumulate with protection from grazing (Cingolani et al. 2005). Thus, continuous grazing in environments with no adaptation to grazing would be detrimental to species diversity (McIntyre and Lavorel 1994; Milchunas et al. 1988). Conversely, non-native plant invader species may have adaptations that better enable them to invade and recover from grazing because they evolved under a grazing regime (Kimball and Schiffman 2003).

Grass species may not be the only taxa affected by livestock grazing. Others studies have indicated an increase in the total density of shrubs (Cipriotti and Aguiar 2012) or increase in shrub cover (Altesor et al. 2006) with grazing. Changes in shrub cover may be due to the palatability of native grasses to livestock, unpalatability of woody species, and subsequent increase in unpalatable woody species (Westoby et al. 1989).

Effects to vegetation in areas that have been excluded from livestock grazing have been studied, but there have been no studies on the SPRNCA that analyze the effects of livestock grazing exclusion on native perennial grass species richness, and invasion of non-native perennial grasses and woody species.

## STUDY AREA DESCRIPTION

The SPRNCA is located in southeastern Arizona approximately 85 km southeast of Tucson, within the Southeastern Arizona Basin and Range Major Land Resource Area. Chihuahuan desertscrub vegetation covers the largest area within the SPRNCA, but other upland vegetation types present within monitoring transects include sacaton grasslands (*Sporobolus wrightii* Munro ex Scribn. and *S. airoides* (Torr.) Torr.), and velvet mesquite (*Prosopis velutina* Wooton) terraces (Makings 2006).

## METHODS

The pace frequency method for rangeland monitoring consists of observing plots along four parallel lines (each five paces apart), with each line containing fifty 40-cm by 40-cm (15-in by 15-in) plots located at one-pace intervals (BLM 1985a). Most pace frequency transects (PFT) were established in 1987 in uplands, with some established in retired agricultural fields to assess effectiveness of restoration. Monitoring of PFTs occurred on an approximate five-year cycle, with monitoring occurring most recently in 2014 to 2015.

Species occurrence, individual species total frequency, and basal or canopy cover frequency of woody species were recorded. A basal (rooted stem) hit is required when recording perennial grasses and woody species. However, if a basal hit is recorded for a woody species, then a canopy hit is not recorded. Canopy is recorded if any live portion of the woody plant occurs over the plot frame. Thus, it is possible to have basal hits of woody species without canopy cover, and vice versa. Total frequency of a woody species includes both basal and canopy hits. Common and scientific names were those currently used by U.S. Department of Agriculture Natural Resources Conservation Service (http://plants.usda.gov/java/).

Significant changes in frequency in perennial grass, half-shrub, shrub, and tree species was analyzed using the methods described by BLM involving confidence intervals (1985b). For half-shrubs, shrubs, and trees, significant changes in basal or canopy cover were determined if significant increases or decreases occurred in total frequency. A two-tailed paired-sample t test was used for normal data, the Wilcoxon paired-sample test was used on data that did not display normality, and the sign test for paired-sample data (Zar 1999) was used to determine significant changes in number of plots with an increase or decrease in a species. The *a priori* significance level was P=0.05.

## RESULTS

### Native perennial grasses

A significant increase in species richness of native perennial grass species was noted (Tables 1 and 2, Figure 1,two-tailed paired-sample t test, t = 6.391, df = 49, P<0.001). There was a significant increase in the number of plots that contained an increase in bristlegrass (*Setaria macrostachya* Kunth and *S. leucopila* (Scribn. & Merr.) K. Schum., sign test for paired-sample, P=0.0005). A significant increase in the number of native perennial grass species were documented in retired agricultural fields (two-tailed paired-sample t test, t = 3.69, df = 8, 0.01<P<0.005).

**Table 1.**
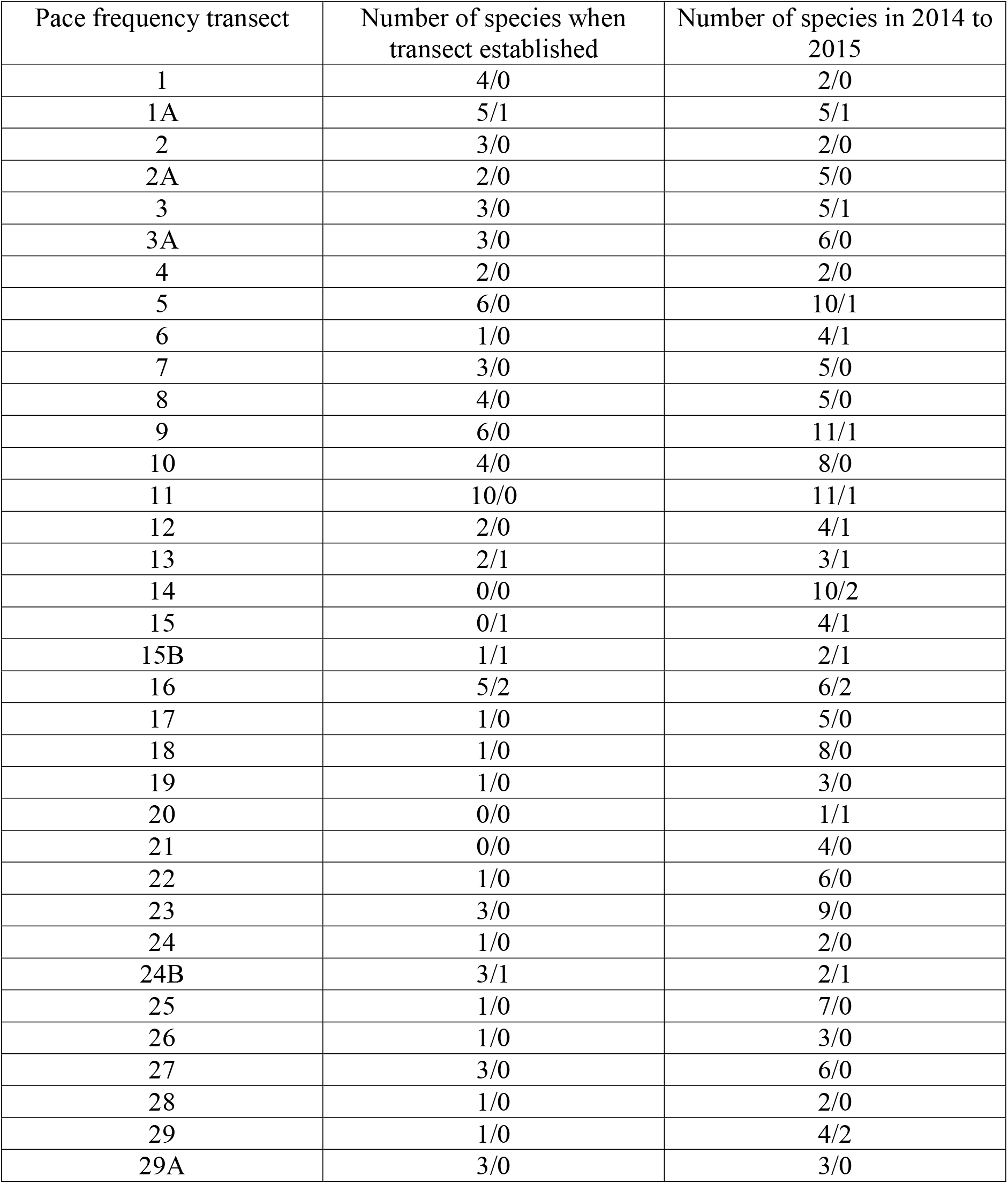
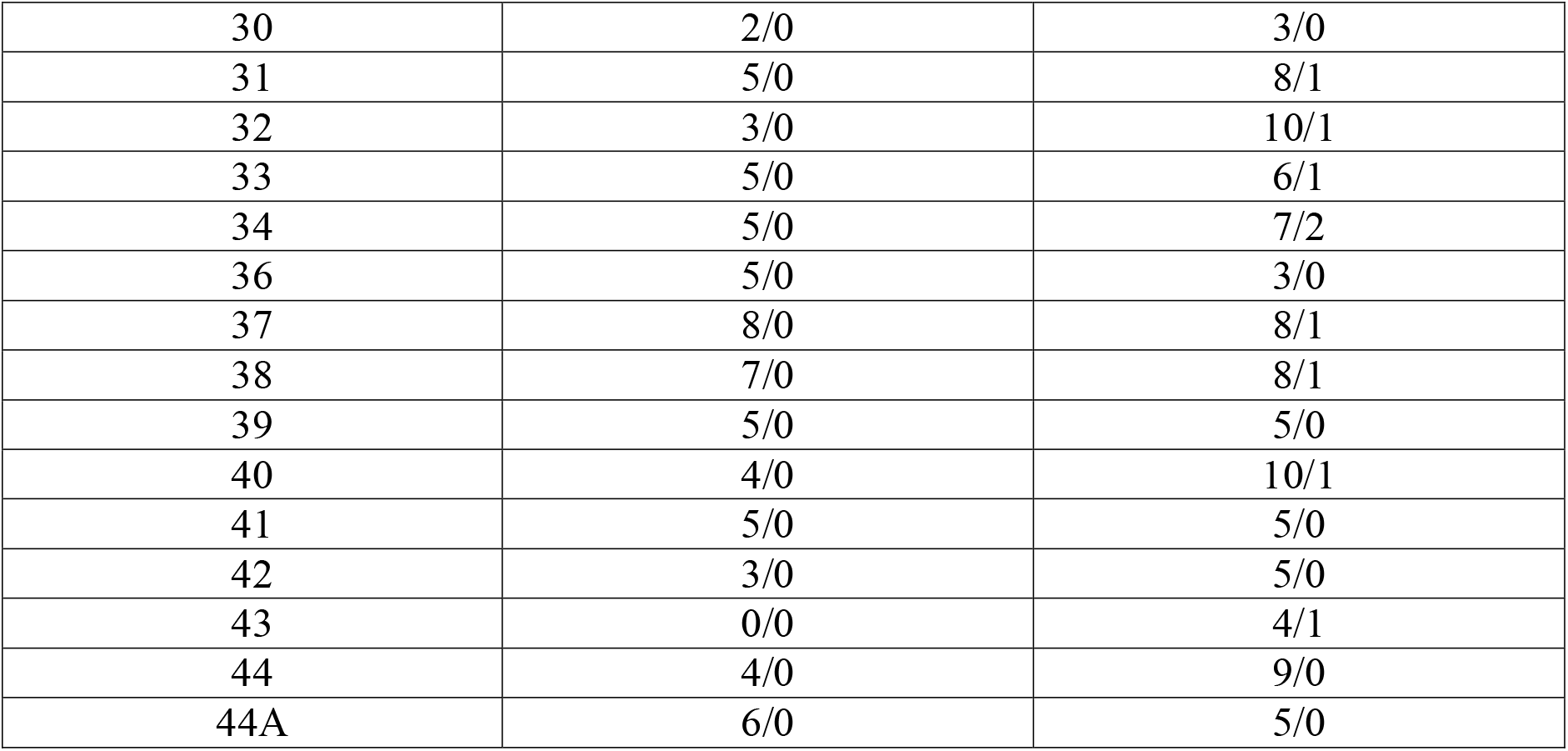
Pace frequency transect with number of native perennial grass species/number of non-native invasive perennial grass species when plot established with number of species in 2014 to 2015, San Pedro Riparian National Conservation Area. For retired agricultural fields (PFT 14, 15, 15B, 16, 17, 18, 19, 20, and 21), 0.01<P<0.005. For all PFT, P<0.001.

| Pace frequency transect | Number of species when transect established | Number of species in 2014 to 2015 |
| --- | --- | --- |
| 1 | 4/0 | 2/0 |
| 1A | 5/1 | 5/1 |
| 2 | 3/0 | 2/0 |
| 2A | 2/0 | 5/0 |
| 3 | 3/0 | 5/1 |
| 3A | 3/0 | 6/0 |
| 4 | 2/0 | 2/0 |
| 5 | 6/0 | 10/1 |
| 6 | 1/0 | 4/1 |
| 7 | 3/0 | 5/0 |
| 8 | 4/0 | 5/0 |
| 9 | 6/0 | 11/1 |
| 10 | 4/0 | 8/0 |
| 11 | 10/0 | 11/1 |
| 12 | 2/0 | 4/1 |
| 13 | 2/1 | 3/1 |
| 14 | 0/0 | 10/2 |
| 15 | 0/1 | 4/1 |
| 15B | 1/1 | 2/1 |
| 16 | 5/2 | 6/2 |
| 17 | 1/0 | 5/0 |
| 18 | 1/0 | 8/0 |
| 19 | 1/0 | 3/0 |
| 20 | 0/0 | 1/1 |
| 21 | 0/0 | 4/0 |
| 22 | 1/0 | 6/0 |
| 23 | 3/0 | 9/0 |
| 24 | 1/0 | 2/0 |
| 24B | 3/1 | 2/1 |
| 25 | 1/0 | 7/0 |
| 26 | 1/0 | 3/0 |
| 27 | 3/0 | 6/0 |
| 28 | 1/0 | 2/0 |
| 29 | 1/0 | 4/2 |
| 29A | 3/0 | 3/0 |
| 30 | 2/0 | 3/0 |
| 31 | 5/0 | 8/1 |
| 32 | 3/0 | 10/1 |
| 33 | 5/0 | 6/1 |
| 34 | 5/0 | 7/2 |
| 36 | 5/0 | 3/0 |
| 37 | 8/0 | 8/1 |
| 38 | 7/0 | 8/1 |
| 39 | 5/0 | 5/0 |
| 40 | 4/0 | 10/1 |
| 41 | 5/0 | 5/0 |
| 42 | 3/0 | 5/0 |
| 43 | 0/0 | 4/1 |
| 44 | 4/0 | 9/0 |
| 44A | 6/0 | 5/0 |

**Table 2.**
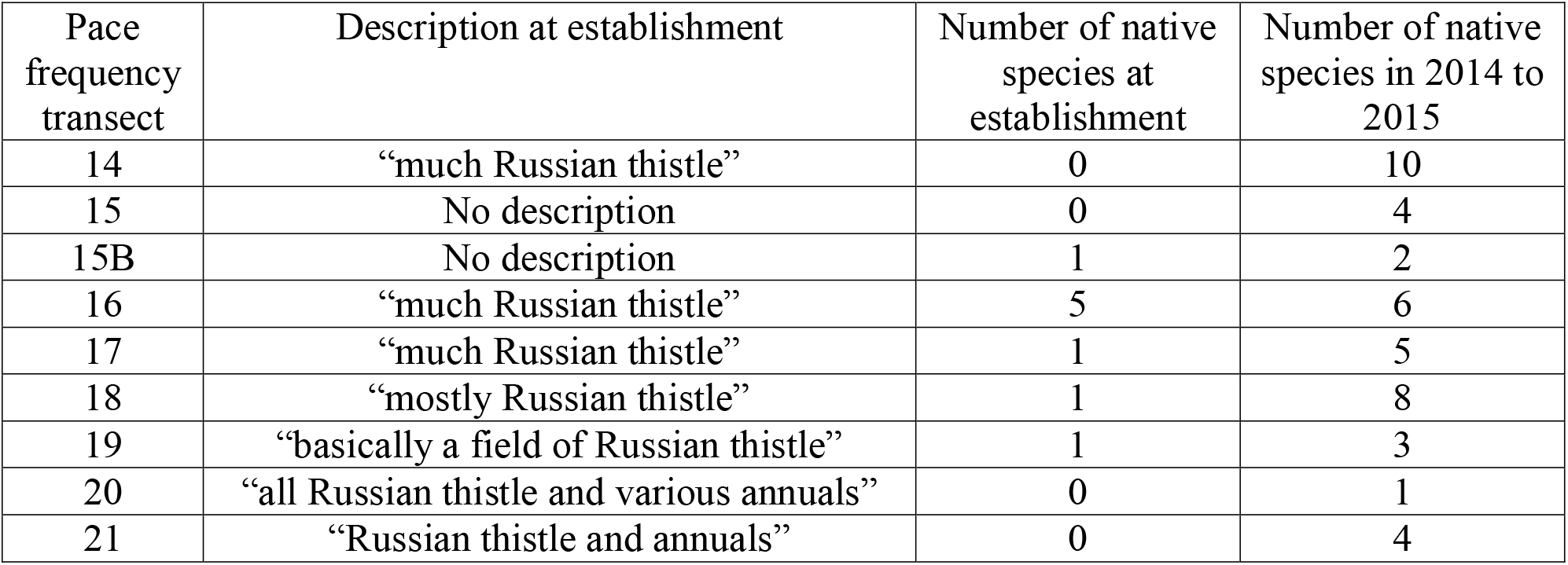
Pace frequency transect in retired agricultural fields with initial qualitative description and number of native perennial grasses, at establishment through 2014 to 2015, 0.01<P<0.005, San Pedro Riparian National Conservation Area.

| Pace frequency transect | Description at establishment | Number of native species at establishment | Number of native species in 2014 to 2015 |
| --- | --- | --- | --- |
| 14 | “much Russian thistle” | 0 | 10 |
| 15 | No description | 0 | 4 |
| 15B | No description | 1 | 2 |
| 16 | “much Russian thistle” | 5 | 6 |
| 17 | “much Russian thistle” | 1 | 5 |
| 18 | “mostly Russian thistle” | 1 | 8 |
| 19 | “basically a field of Russian thistle” | 1 | 3 |
| 20 | “all Russian thistle and various annuals” | 0 | 1 |
| 21 | “Russian thistle and annuals” | 0 | 4 |

**Figure 1.**
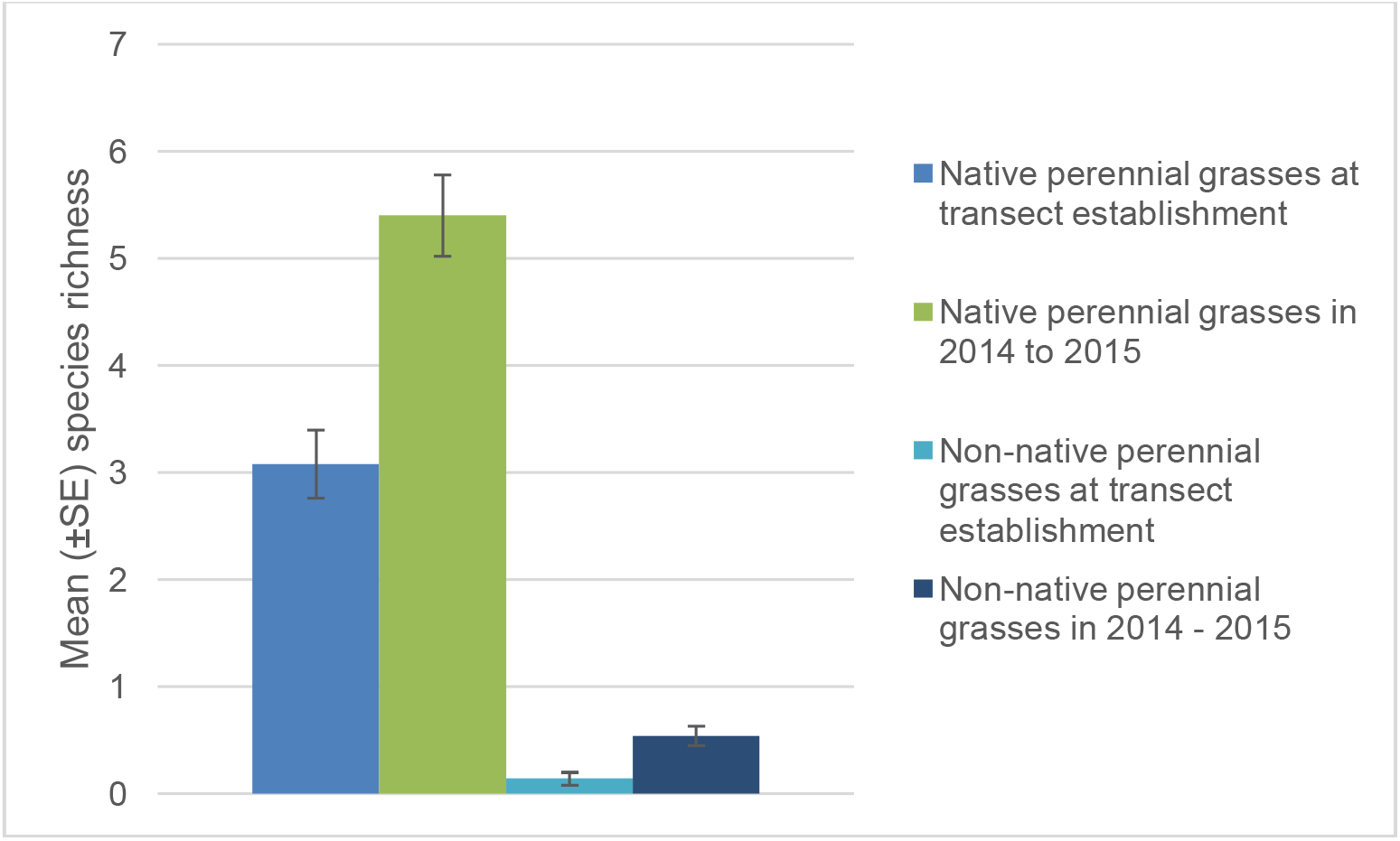
Mean (± standard error) species richness of native perennial grasses and non-native perennial grasses at transect establishment and in 2014 to 2015, San Pedro Riparian National Conservation Area. Species richness of native perennial grasses increased significantly (P<0.001). Species richness of non-native perennial grasses did not significantly change (0.10<P<0.20).

Although not within the significance level, most significant increases of native perennial grasses included green sprangletop (*Leptochloa dubia* (Kunth) Nees, P=0.06), and cane bluestem (*Bothriochloa barbinodis* (Lag.) Herter, P=0.06). No other significant changes in native perennial grass frequency were observed.

### Non-native perennial grasses

Lehmann lovegrass (*Eragrostis lehmanniana* Nees) and Johnsongrass (*Sorghum halepense* (L.) Pers.) remained the same two species present prior to grazing exclusion and in 2014 to 2015. There was no significant change in species richness of non-native perennial grasses (Table 1, Figure 1, Wilcoxon paired-sample test, T_=479, n =50, 0.10<P<0.20).

However, Lehmann lovegrass had a significant increase in the number of plots that contained this species (sign test for paired-sample, P=0.03126), but Johnsongrass did not significantly increase in the number of plots with this species (sign test for paired-sample, P=1.0).

### Woody species

Velvet mesquite exhibited a significant increase in total frequency (sign test for paired-sample, P=0.001). However, velvet mesquite did not exhibit a significant increase or decrease in basal hits (sign test for paired-sample, P=0.62), but did have a significant increase in canopy cover only (test for paired-sample, P=0.00002). There was a significant decrease in total burroweed (*Isocoma tenuisecta* Greene, sign test for paired-sample, P=0.04), and basal zinnia (sign test for paired-sample, P=0.04). Although not within the significance level, the next most significant increase included total frequency (i.e. basal and canopy) of mariola (*Parthenium incanum* Kunth, P= 0.06), No other significant changes in half-shrub, increaser half-shrub, shrub, tree, or succulent frequency were determined.

## DISCUSSION

### Native perennial grasses

On the SPRNCA, results of this study support the hypothesis that native perennial grass species richness increases with livestock grazing exclusion. Similarly, on nearby Las Cienegas National Conservation Area (LCNCA), ungrazed key areas had increases in perennial grass richness compared to adjacent grazed areas (Gori and Schussman 2005). American bison (*Bison bison*) are not native to Arizona (Reynolds et al. 2003). The increase in native perennial grass richness on SPRNCA with grazing exclusion may illustrate at least a short-term lack of adaptation of local native perennial grasses because this flora has not evolved under this disturbance regime.

The frequency of bristlegrass significantly increased along with a significant increase in native perennial grass richness. Perhaps bristlegrass is an indicator species for restoration, and indicates conditions are improving enough so that other species will follow. As shown by nearly significant increases in green sprangletop and cane bluestem, additional time of grazing exclusion may show additional increases in richness and frequency of native perennial grasses. Some studies of long-term grazing exclosures suggested very slow rates of vegetation recovery in arid and semi-arid environments (Valone et al. 2002), and recovery of the native perennial grass component on SPRNCA may still be in the earliest stages. It is noteworthy that perennial grass species richness has increased with grazing exclusion, even with ongoing long-term drought.

Likewise, initial monitoring efforts noted that the retired agricultural fields were largely Russian thistle with few other species at transect establishment. Since those observations, a significant increase in species richness of native perennial grasses in the retired agricultural fields was documented during this study. Although not quantified, native perennial grasses with increases in the agricultural fields included green sprangletop, giant sacaton, large-spike bristlegrass, purple threeawn, sideoats grama, and cane bluestem. Continued passive restoration of the retired agricultural fields with native perennial grasses may also allow improved water infiltration for maintenance and restoration of the riparian area.

The SPRNCA has large areas of upland habitat, which continues to be degraded due to ongoing topsoil loss and erosion. Grazing may increase soil disturbance and erosion (Greene et al. 1994), which favors unpalatable perennial shrubs (Westoby et al. 1989). Shrublands offer less surface protection from erosion than historic grassland vegetation (Zeedyk and Clothier 2009), but healthy grasslands better allow infiltration of precipitation than shrublands (Deboodt 2009). The resulting improved water infiltration from healthy grasslands may provide water to the aquifer needed for maintenance, enhancement, and restoration of the riparian area.

### Non-native perennial grasses

A significant increase in the number of plots with exotic Lehmann lovegrass occurred with grazing exclusion on SPRNCA, but there was not a significant increase in frequency on plots that were already invaded. These results indicate Lehmann lovegrass invaded additional sites, but did not change in frequency with grazing exclusion in the sites that were originally invaded. Similarly, in another study, Lehmann lovegrass density increased with time and was not affected by grazing exclosure (McClaran and Anable 1992). In yet another study, there was some weak evidence that heavy grazing favored invasion by exotic plant species, with negative effects of grazing on native plant species richness at broader landscape scales (Dorrough et al. 2007).

Results of this study indicate that Lehmann lovegrass expansion occurs in previously uninvaded areas even without livestock grazing, likely because the ecosystem has already been profoundly altered over long periods of time by historic land management and subsequent ongoing soil erosion.

On the SPRNCA, approximately 10,000 ha (24,710 ac) of upland areas have soil erosion, where active sheet flow, rills, and gullies occur. Approximately 1,000 ha (2,471 ac) occurs on upland soils rated “fragile” by the U.S. Department of Agriculture Natural Resources Conservation Service soil web survey. This rating is based on soils that are weakly structured, have low organic material, occur in a dry climate, and may have slopes. The current degraded state of soil horizons may have influenced the subsequent invasion of Lehmann lovegrass to new sites on SPRNCA.

### Woody species

Significant changes in frequency of velvet mesquite on SPRNCA were the result of increased canopy cover, and not basal hits, indicating the same number of rooted plants existed prior to and after grazing exclusion. These same rooted basal plants have merely grown and increased in canopy cover during the livestock grazing moratorium. Cattle readily consume and disperse viable mesquite seeds (Kramp et al. 1998), passage of seeds through the ruminant gut may enhance seed germination (Scifres and Brock 1969, Wilson et al. 2001), and seeds in cattle dung have higher seedling survival rates compared to areas without cattle dung (Brown and Archer 1987). Consequently, encroachment and reinvasion of mesquite into grasslands may be accelerated by livestock consuming and dispersing seeds (Gibbens et al. 1992, Lerner and Peinetti 1996). In another study, mesquite seedlings occurred in high density on areas with cattle, in contrast to an absence of seedlings on areas without cattle, suggesting that rates of invasion of grasslands by mesquite would have increased following the introduction of domestic ungulates (Brown and Archer 1988). Anecdotal evidence of subsequent mesquite invasion is qualitatively observed on nearby LCNCA, where grazing is allowed, with areas cleared of mesquite now invaded with small mesquite seedlings. Thus, past cattle grazing was likely a primary factor leading to the wide distribution and density of velvet mesquite on SPRNCA as evidenced by the lack of significant increase in basal hits with grazing exclusion.

Burroweed is another woody species that increases under grazing pressure. Low palatability to livestock, and reduced frequency of fire in desert grasslands, have generally increased the abundance of this species (Cronin et al. 1978). However, results of this study indicate that total burroweed frequency has decreased on SPRNCA, even with fire suppression but with grazing exclusion. This study’s results are similar to Schmutz and Smith (1976), who found a slight decline in frequency of burroweed on an area protected from grazing. Conversely, zinnia is short-lived (Goldberg and Turner 1986), and sensitive to drought. The significant decline in zinnia may anecdotally illustrate the effects of long-term drought on SPRNCA and not effects of grazing exclusion.

## MANAGEMENT IMPLICATIONS

Determining the extent of degradation in semi-arid rangelands will always be hampered without reference sites that have never been subject to grazing (Fensham 2011), or without reference sites that have long been excluded from grazing. Livestock grazing should continue to be excluded on the SPRNCA in order to provide a rare opportunity to scientifically compare the effects of exclusion to other grazing regimes on annual and perennial native grasses, non-native grasses, shrubs, and trees, especially because an increase in species richness and frequency of native perennial grasses has been documented on the SPRINCA during this study.

Additionally, continued grazing exclusion on SPRNCA may allow future increases in plant richness, which is one of many complex and interconnected factors determining species richness in animal taxa (Hawkins and Pausas 2004; Wolters et al. 2006). Native perennial and annual grasses and forbs may be important browse and pollinator species for many species of wildlife. Increased plant species richness on SPRNCA may subsequently improve animal richness.

Monitoring of the PFTs on the SPRNCA should continue and include annual and perennial forbs that are important as pollinator species and as browse for wildlife.

Ongoing soil erosion on SPRNCA occurs in upland wildlife habitat and, for effective management of wildlife habitat, monitoring of erosion and erosion control activities should occur.

Full wildfire suppression has been the objective since the inception of SPRNCA, even in upland habitats with woody species invasion. Wildfire in the area historically occurred primarily during late spring and early summer from lightning starts. Thus, prescribed fire should be used as the preferred tool to control woody species in upland vegetation on SPRNCA. Grazing exclusion would be necessary to provide fine fuels (grasses and forbs) for effective fire treatments. Unlike broadcast herbicide treatment to control woody species, the use of prescribed fire leaves a mosaic of habitat and allows for survival of other dicot species important for pollinators and other wildlife.

## Notes

### Competing Interest Statement

The authors have declared no competing interest.

